# Genome-Wide Exploration of the Opportunistic *Providencia stuartii* Unveils the Novel Genetic Interactions with the Virulence Gene of Diarrheal Pathogens

**DOI:** 10.1101/2023.11.22.568233

**Authors:** Mohammad Uzzal Hossain, Md. Shahadat Hossain, A.B.Z Naimur Rahman, Shajib Dey, Zeshan Mahmud Chowdhury, Arittra Bhattacharjee, Ishtiaque Ahammad, Md. Abdul Aziz, Abu Hashem, Keshob Chandra Das, Chaman Ara Keya, Md. Salimullah

## Abstract

Diarrhea typically indicates an intestinal disorder, which can occur from viruses, parasites or bacterial infection. Along with the common diarrhea-causing pathogens, opportunistic bacteria may also play a role in the etiology of diarrheal disease. One of the opportunist’s bacteria that can cause diarrhea in both children and adults is *Providencia stuartii*. Therefore, the goal of this study is to explore the genetic mechanism of the opportunistic *P. stuartii* in microbial interactions with common diarrheal pathogens. Hence, *P. stuartii* was identified by utilizing the morphological observation and molecular techniques. Afterwards, the entire genome of *P. stuartii* was sequenced, assembled and annotated to explore the genomic insights. In addition, the virulence genes of 100 whole genome sequences from ten prevalent diarrhea-causing bacteria were identified and prioritized. Finally, the system biology approach was used to predict the protein-protein interaction network between *P. stuartii* and the virulence genes. The results of the present study suggests that complete genome sequencing of this bacteria contains 4011 proteins, which are crucial for this bacterium to survive. Additionally, 16 gene clusters provide 207 interacting genes that could interact with biological and molecular function, subcellular localization and pathway. The microbial interaction accompanying the virulence gene was found in all 10 diarrhea-causing bacteria except *Clostridium difficile*. These findings of this study could aid in the exploration of *Providencia stuartii* as the major causative agent of diarrhea. Additionally, the pathophysiology of diarrhea can be investigated using the microbial interactions between *P. stuartii* and the typical diarrheal bacteria. The results of this study may therefore be used to determine the most effective therapeutic targets for the development of medications to treat diarrhea.

## 1. Introduction

Diarrheal disease constitutes a significant global public health challenge, contributing to substantial rates of morbidity and mortality. It stands as one of the primary reasons for outpatient consultations, hospital admissions, and the worldwide burden of years of life lost (YLL) across all age groups [1]. Roughly 1.6 million deaths transpire annually on a global scale as a result of diarrhea, with the most profound impact felt in developing nations and economically marginalized regions [2].

Globally, diarrhea contributed to 15% among children under the age of five [3,4]. Out of all child deaths from diarrhea, 78% occur in the Southeast Asian and African regions [5–7]. By the year 2030, projection estimated that infectious diseases will cause around 4.4 million child deaths under the age of five annually [8]. The disease is particularly dangerous for young children, who are more susceptible to dehydration and nutritional losses, especially those living in low and middle incomed countries. The accurate extent of diarrheal disease impact within vulnerable populations in Bangladesh remains inadequately assessed, and only a few studies were conducted to examine the healthcare seeking behavior for diarrhea, which has crucial implications on the outcomes of diarrheal diseases [9,10].

Antibiotic is mostly given for the treatment with bacterial diarrhea. But the high failure rate is increasing day by day due to acquired resistance to commonly used antibiotics [11]. Most diarrheal pathogens have developed resistance against the major classes of antibiotics commonly used for mitigating diarrheal symptoms [12]. However, the number of infections caused by multidrug-resistant bacteria is increasing globally and the prospect of untreatable infections is becoming a reality [13]. The most frequently identified pathogenic organisms causing bacterial diarrhea are *Escherichia coli*, *Shigella, Salmonella*, *Campylobacter* and *Clostridium spp*. [14,15]. The existing approaches to addressing diarrhea and its impact on both adults and children fall short of achieving complete effectiveness. Recent researches are focused on handling the disease with oral rehydration, zinc supplementation, vaccination, antibiotics, prebiotics and probiotics [16–19].

However, very few effective research is ongoing focusing on the genomics of the diarrhea causing pathogens for targeted interventions and microbial interaction underlying the emergence and recurrence of the disease. Microbial interactions among the frequently documented pathogens causing diarrhea are frequently observed in connection with typical bacteria such as *Salmonella, Campylobacter jejuni, Shigella, Listeria monocytogenes, Vibrio parahaemolyticus, Vibrio cholerae* as well as different types of pathogenic *Escherichia coli* including Enterohemorrhagic *Escherichia coli* (EHEC), Enteropathogenic *Escherichia coli* (EPEC), Enteroinvasive *Escherichia coli* (EIEC), Enterotoxigenic *Escherichia coli* (ETEC), and Enteroaggregative *Escherichia coli* (EAEC) [15,20,21].

In addition to the commonly associated bacteria, various opportunistic bacteria might contribute to the development of diarrhea. *Providencia stuartii* (*P. stuartii*) is a rare opportunistic organism that causes healthcare-associated infections, such as acute enteric infection, urinary tract infection, and lung diseases [4]. The organism is typically isolated from human secretions, including urine, sputum, blood, stool, and wound cultures [22]. Alongside with other infection and diseases, *Providencia spp.* notably strains of *P. stuartii* was isolated from the pool of 130 patients with travelers’ diarrhea [23]. In certain studies, *Providencia* was identified as a causative agent of diarrhea, supported by the clinical spectrum indicating its role in causing invasive diarrheal conditions [24,25]. *P. stuartii* was first isolated in 1904 by Rettger, and was named in 1951 by Kauffmann [25]. Following its emergence, *P. stuartii* led to numerous outbreaks documented in Brazil, China, Taiwan and Tunisia. Predominantly reported cases were within hospital facilities, particularly in intensive care units (ICUs), long-term care facilities (LTCFs), and burn units [24–30].

According to recent studies, opportunistic *P. stuartii* were reported as carbapenemases antibiotic resistant and significant threat to public health and associated risks [4,32]. However, the complete genomics of the *P. stuartii* is not extensively studied that might provide insights of the essential genes, gene activity, protein interactions and genetic variations. This study aims to explore the genomics variation and the essential protein interactions of *P. stuartii* with bacterial diarrhea pathogens. Utilization of Next-generation sequencing (NGS) and comprehensive genome analysis for *P. stuartii* is promising [33]. These approaches might unveil microbe-microbe interactions with other diarrhea-causing bacteria, offering insights into why this opportunistic pathogen may have gained dominance and become a potential contributor to diarrhea and associated disease. Research on *P. stuartii*, adapting the synergy of system biology and computational algorithms holds promising prospects for advancing diarrheal therapeutics, targeted therapeutics and novel antibiotics for tailored therapeutic interventions.

## 2. Materials and Methods

### A. Wet Laboratory

### 2.1 Clinical sample collection

Eleven fecal samples were collected from two hospital facility in the metropolitan city Dhaka, Bangladesh (N 23° 46.387980 E 90° 22.168260 and N 23° 42.679980 E 90° 24.065940) and selected based on the children diarrhea patient treatment facility. Sterile stool collection tubes and sterile cotton swabs were used to collect fecal samples and inoculated in the fresh Tryptic Soy Broth (TSB) media. The samples were immediately transported to storage facility and were incubated at the incubator at 37°C. **Supplementary Table 1** contains sample metadata, providing information on patients’ age, origin and gender.

### 2.2 Bacterial culture and DNA extraction

The microorganisms were initially incubated for 48 hours in Triptic Soy Broth (TSB) until the media turned opaque. OXOID S.S. AGAR and MacConkey Lactose Agar were used as medium for a 24-hour growth period to identify enteric bacteria through colony characteristics. *Providencia* like colonies were then isolated, with single colonies transferred to MacConkey Lactose Agar for further characterization. Pure cultures were developed on Nutrient agar for 24 hours. The genomic DNA from each of the four bacterial pure colony samples was extracted using the Thermo Scientific GeneJET Genomic DNA Purification Kit following the manufacturer’s instructions. Concentration and purity were estimated by Thermo Scientific NanoDrop 2000/2000c and stored in the −20°C freezer.

### 2.3 16S rRNA universal primer-based amplification and cycle sequencing

Amplicon generation and library preparation were performed using the forward primer 27F (5’-AGAGTTTGATCMTGGCTCAG-3’) and reverse primer 1492R (5’-GGTTACCTTGTTACGACTT-3’), targeting the V5-V9 regions of the 16S rRNA gene. The PCR was conducted with a reaction volume of 15 μL, comprising 7.5 μL of GoTaq® Master Mix, 0.75 μL each of the forward and reverse primers, 3 ng of template DNA, maintaining a ratio of 100 ng/μL for the template DNA, and 3 μL of nuclease-free water to reach the final reaction volume. The amplified PCR product was gel-documented, and the PCR band was visualized using the gel-documentation system. The generated amplicons were checked for concentration, and purity was estimated using the Thermo Scientific NanoDrop 2000/2000c. Subsequently, cycle sequencing was performed using the Veriti 96 well Thermal Cycler from Applied Biosystems by Thermofisher Scientific® with a 10 μL reaction volume for 25 cycles of sequencing.

### 2.4 Sanger sequencing and phylogenetic analysis

Sanger sequencing was performed using the 3500 Genetic Analyzer from Applied Biosystems by Thermo Fisher Scientific. The purified samples were loaded onto a 96-well plate and sequenced using the analyzer. Sanger sequence reads retrieved from the sequencing was subjected to filtration and the reads with low confidences were removed and aligned using the local alignment tool MEGA [34]. Afterwards, the phylogenetic tree was constructed using the query sequence reads and fourteen sequences retrieved from NCBI and the phylogenetic relatedness of the clinical isolates species was determined using the MEGA [34].

### B. Next generation sequencing

### 2.5 Next generation sequencing (NGS), assembly and annotation of the clinical isolate

The Sanger sequences that identified and confirmed clinical isolate DNA were sequenced using the MiSeq™ System by the Illumina sequencing platform. Data preprocessing and quality control were performed using the Linux terminal and FASTQ files were analyzed using the FASTQC tool, and the adapter sequence was trimmed using the Trimmomatic tool [35]. Assembly was conducted using Unicycler, following the pipeline for prokaryotic genome assembly. Prokka and RAST were implemented for the annotation of the contigs generated by the hybrid assembler [36,37]. Using the DNAPlotter application the whole genome of *P. stuartii* circular view was generated which plotted the contigs and GC content confidence in graphical view [38]. Proteins were identified using the awk function in Linux terminal command lines, and the protein sequences of the essential proteins were filtered.

### 2.6 Retrieval of hundred whole genome sequence of diarrheal pathogens

Publicly available dataset from the NCBI web specific SRA run selector was used to retrieve the hundred whole genome sequences from ten microbial organisms namely *S. dysenteriae, S. flexneri, Salmonella enterica* Typhimurium*, Salmonella enterica* Enteriditis, *V. cholerae, V. parahemolyticus, C. difficile, C. jejuni*, two strains of *E. coli* i.e., Enteroinvasive *Escherichia coli* (EIEC) and Enterotoxigenic *Escherichia coli* (ETEC) and from each 10 SRA data was downloaded using the SRA-toolkit.

The 100 whole genome data have been retrieved from Asia continent and 5 different countries including Bangladesh (**Supplementary Table 2**). Data preprocessing and quality control was performed as the FASTQ file underwent analysis using the FASTQC tool, and the adapter sequence was subsequently trimmed using the Trimmomatic tool [35].

### 2.7 Assembly and annotation of hundred diarrheal pathogen whole genome

The assembly was carried out for 100 bacterial whole genomes, involving 10 bacterial strains using hybrid de novo assembler Unicycler [39]. The annotation of the one hundred bacterial genome assembly was performed using the Prokka [36]. The annotated files of 100 bacterial genome proteins were analyzed, the essential proteins from the ten bacterial strains were separately retrieved, hypothetical proteins were excluded. The annotation process provided the characterized and the hypothetical protein sequences in different file formats.

### C. Systems biology approach

### 2.8 Virulence gene identification of diarrheal pathogens

Virulence genes from the 10 pathogenic organisms were screened using the Virulence factor database (VFDB) database for virulence factor identification [40]. The Abricate (version 0.8.13) tool was used to identify the genes that predicted as virulent and the VFDB database was utilized by this tool.

### 2.9 PPI interaction between virulence genes and essential proteins of *Providencia stuartii*

The protein sequences were employed to establish a protein-protein interaction network, focusing on the essential genes from the clinical isolate *P. stuartii* in conjunction with the virulence factors (VF) found in 100 diarrheal whole genome datasets. The STRING server was utilized to identify the interactions among these proteins [41]. This process was repeated to create protein-protein interaction networks involving the virulence genes and essential proteins for each bacterium.

### 2.10 Identifications of interacting partners among virulence genes and *Providencia stuartii*

The network of diarrheal bacteria and *Providencia stuartii* was analyzed to detect highly connected proteins based on the degree distribution of nodes. The Cytoscape tool plugin Cytohubba was utilized to identify the top 30 nodes with the highest degree distribution between interacting proteins [42]. The top 30 highly interacting proteins and clustered into one or two groups using the K-means clustering method through the STRING web server [41].

The STRING-generated files were visualized using the Cytoscape tool, and interaction analysis was performed predicting associated nodes. The gene ontology of biological function, molecular process, and the KEGG pathways was identified using the prediction model of STRING. The cluster networks were visualized through different layout options, and high-quality network images were generated for more detailed visualization.

## 3. Results

### A. Wet Laboratory Experiments

### 3.1 Identification of opportunistic *P. stuartii* pathogen in diarrheal patients

Gram-negative bacteria cultured and sub-cultured in enteric culture media lead to the identification of four samples exhibiting colony properties associated with opportunistic pathogens. *Providencia* like bacteria colonies cultures showed the characteristic opaque colony in SS Agar and McConkey Agar media **(Figure 1)**. Subsequently, genomic DNA extracted from these four distinct pure bacterial colonies and amplified PCR products presented bright bands around the 1400 bp region confirming the presence of bacterial DNA in four samples **(Figure 2)**. Sanger sequenced reads were analyzed with NCBI blastn suite and phylogenetic tree established from the sequenced clinical isolate and other 14 sequences showed that one query sequence cluster into a branch of *P. stuartii* which established the confidence of the blastn prediction to be accurate **(Figure 2)**. The presence of this uncommon opportunistic pathogen, *P. stuartii* was detected in the fecal sample of diarrheal patients.

**Figure 1:**
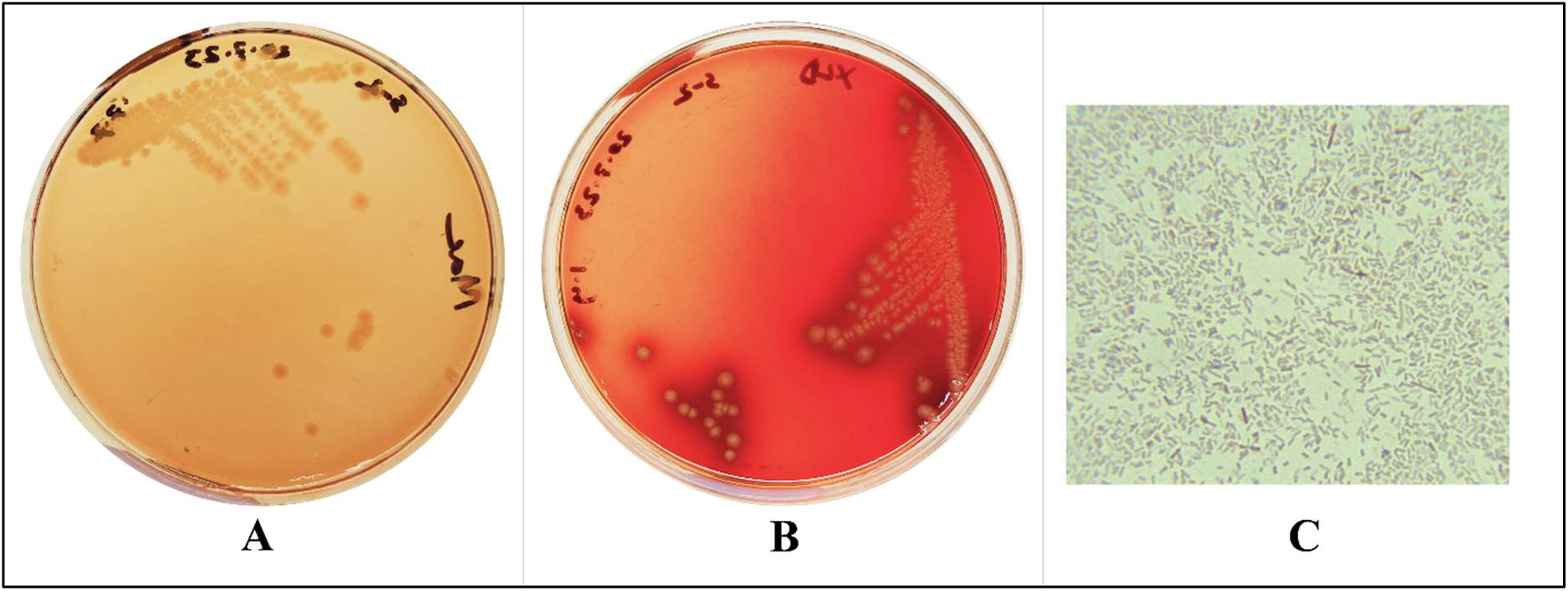
Bacterial colony of the opportunistic pathogen *P. stuartii*. Colonies of *Providencia stuartii* were non-lactose fermenting, orange to red-colored on (A) MacConkey agar and yellow colonies with acid production that changes the pH of the medium on (B) XLD agar plates. Microscopic features (C) of *P. stuartii* Gram staining results were red and appeared as short rods, which indicating that the strain was gram negative.

**Figure 2:**
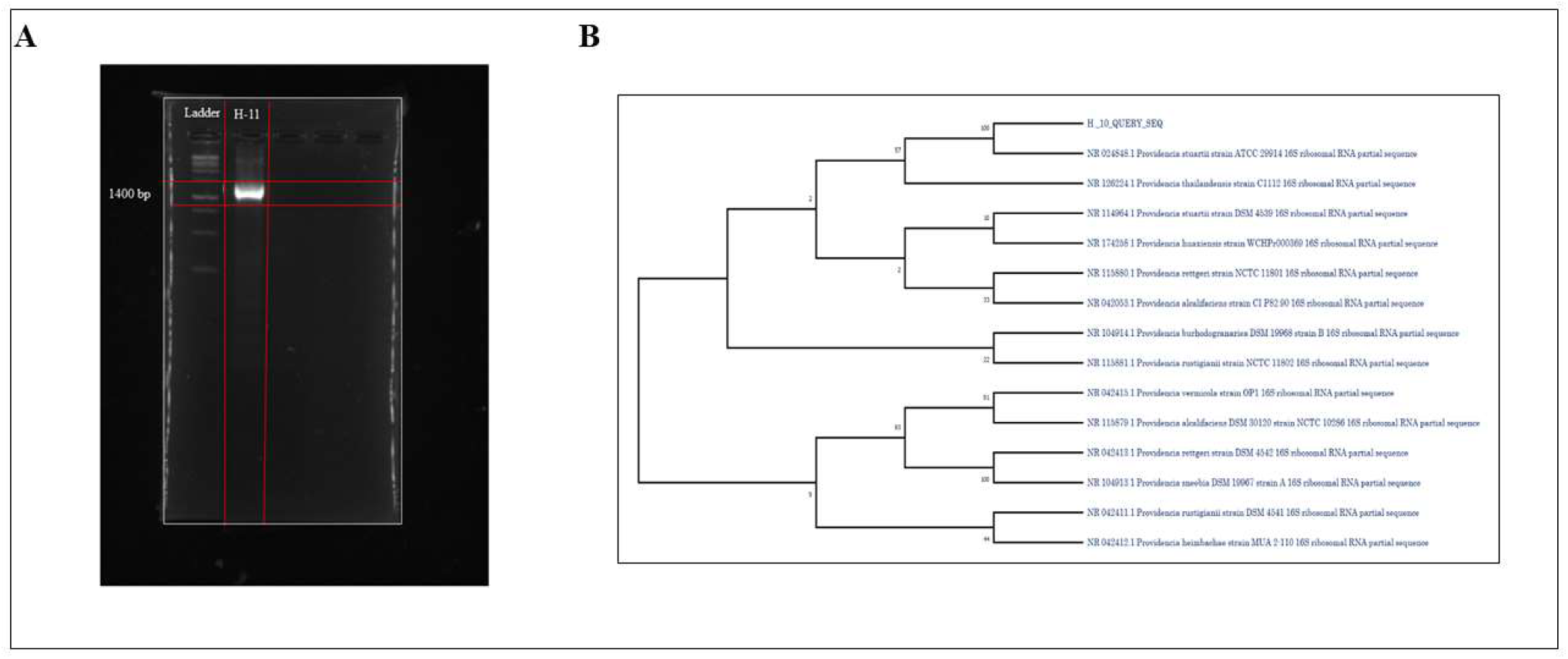
Gel documentation of PCR bands and Phylogenetic tree of *Providencia* sequences. Four specific colony selected for sanger sequencing PCR products formed a deep band around 1400bp specific for 16s rRNA confirming the presence of high-quality genomic DNA (**A**). The query sanger sequence reads of our study denoted by H_10_QUERY_SEQ sample and the 14 sequences are retrieved from NCBI, sequences show the cluster with *Providencia stuartii* strain ATCC 029914 with high similarity (**B**). Annotation of the *P. stuartii* predicted that the sequence contains 159 contigs, a total of 4354629 bases, 4011 coding sequences, 71 tRNAs and 1 rRNA in *Providencia stuartii* bacteria whole genome. Quality Assessment Tool for Genome Assemblies (QUAST) report depicted that the assembly have total length of 4339880 bp, with the largest contig of 784289 bp (**Supplementary Figure S1**).

### B. Next Generation Sequencing Analysis

### 3.2 Genomic features of *Providencia stuartii*

Hybrid assembler Unicycler assembled the paired end reads of the *P. stuartii* sequence and generated assembly files that contained 159 contigs. The GC content was predicted 41.29% which is excellent and N50 value of 259,815 base pairs suggesting that half of the assembled sequences in the genome are 259,815 base pairs or longer (**Figure 3)**.

**Figure 3:**
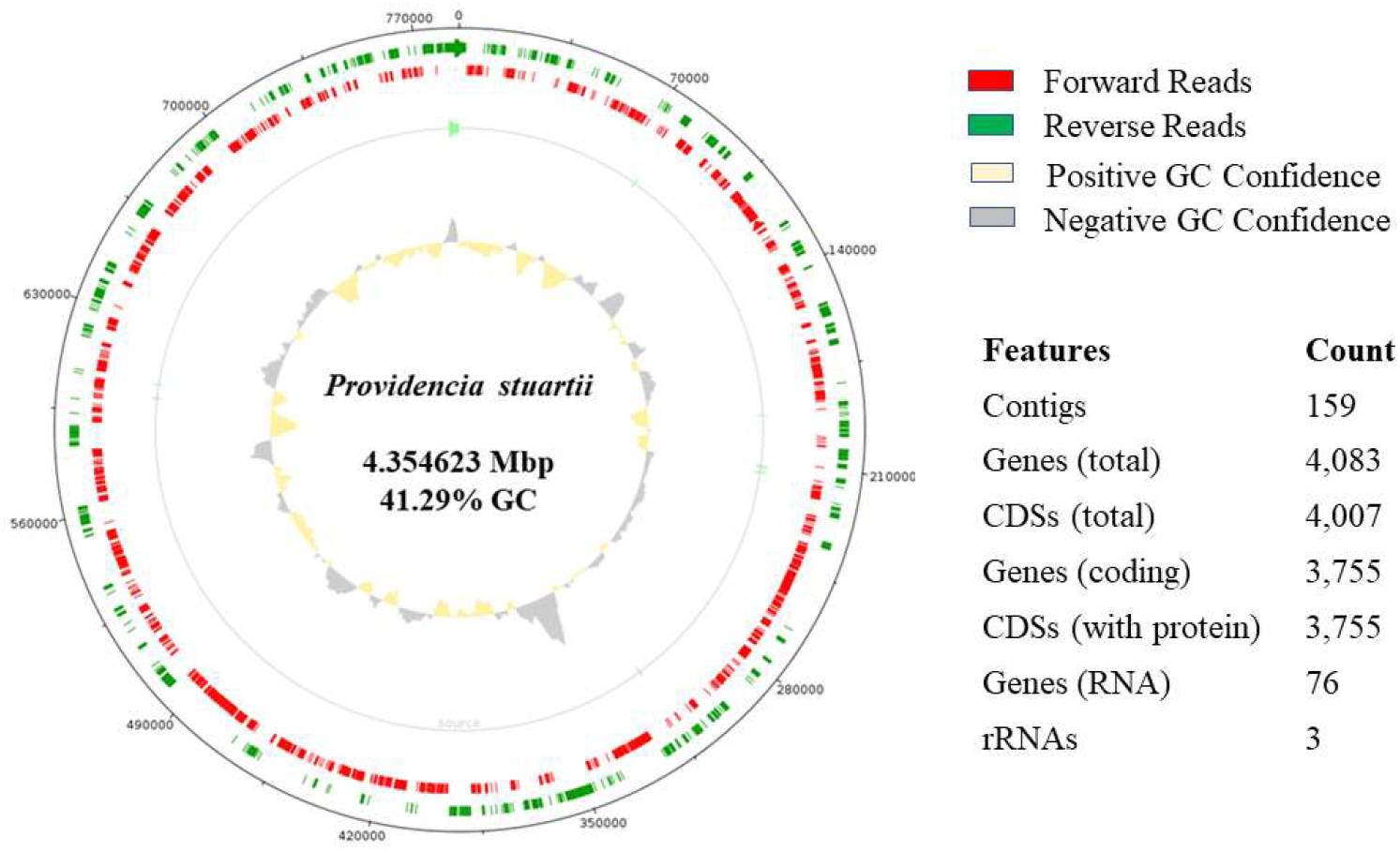
Whole genome plot of *Providencia stuartii*. Green line represents the forward reads and the red line reverse reads of the whole genome assembly, grey and yellow colors represent the confidence of the GC value; consisting of 159 contigs, totaling of 4083 genes and GC value of 41.29%.

### 3.3 Virulence Genes of Diarrheal Pathogen

The Virulence Factor Database (VFDB) for pathogenic bacteria has identified and predicted a varied number of virulence genes across different bacterial species. Notably, *S. dysenteriae* contains 87 virulence genes, *S. flexneri* with 84, *S. Typhimurium* with 152, *S. enteritidis* with 106, *V. cholerae* with the highest count of 159, followed by *V. parahemolyticus* with 46, and *C. difficile* with the lowest count of 10 virulence genes. Additionally, EIEC and ETEC are noted to possess 86 and 82 virulence genes respectively (**Supplementary Table 3)**.

### C. Systems Biology Analysis

### 4.7 Highly connected genes of diarrheal pathogens

*Providencia stuartii* yielded a comprehensive network of 1934 nodes and 10477 edges (**Supplementary Figure S2)**. The virulence gene network of the ten bacterial pathogens and the complete *P. stuartii* network were analyzed and top 30 nodes with the highest number of degree distribution predicted were used for the protein-protein interaction.

### 4.8 Novel insights of interacting pathogens

The network involving *Shigella dysenteriae* and *P. stuartii* formed single cluster encompassing 38 genes, featuring three virulence genes namely ompA, fes and rpoS. Within the highly interacting protein network of *Providencia stuartii*, the *Shigella dysenteriae* cluster revealed the presence of 33 essential genes primarily 27 ribosomal proteins rplO, rplP, rpsE, rpsJ, rpsN, rpsP, rpsQ, rpsM, rpsF, rpsE, rpsV, rpsD, rpsK, rpsC, rplK, rplR, rpsB, rpsN, rpsQ, rplW, rplC, rpaS, rpoA, rplB, rplX, rplA, rpmD. Additionally, transcription factors rpoC, rpoB, rpoA and few other proteins rspH, frr, guaA, tsf were visualized. Functionally, this cluster biological processes are associated with translation and subcellular localization around polysomal ribosomes (**Figure 4A**).

**Figure 4:**
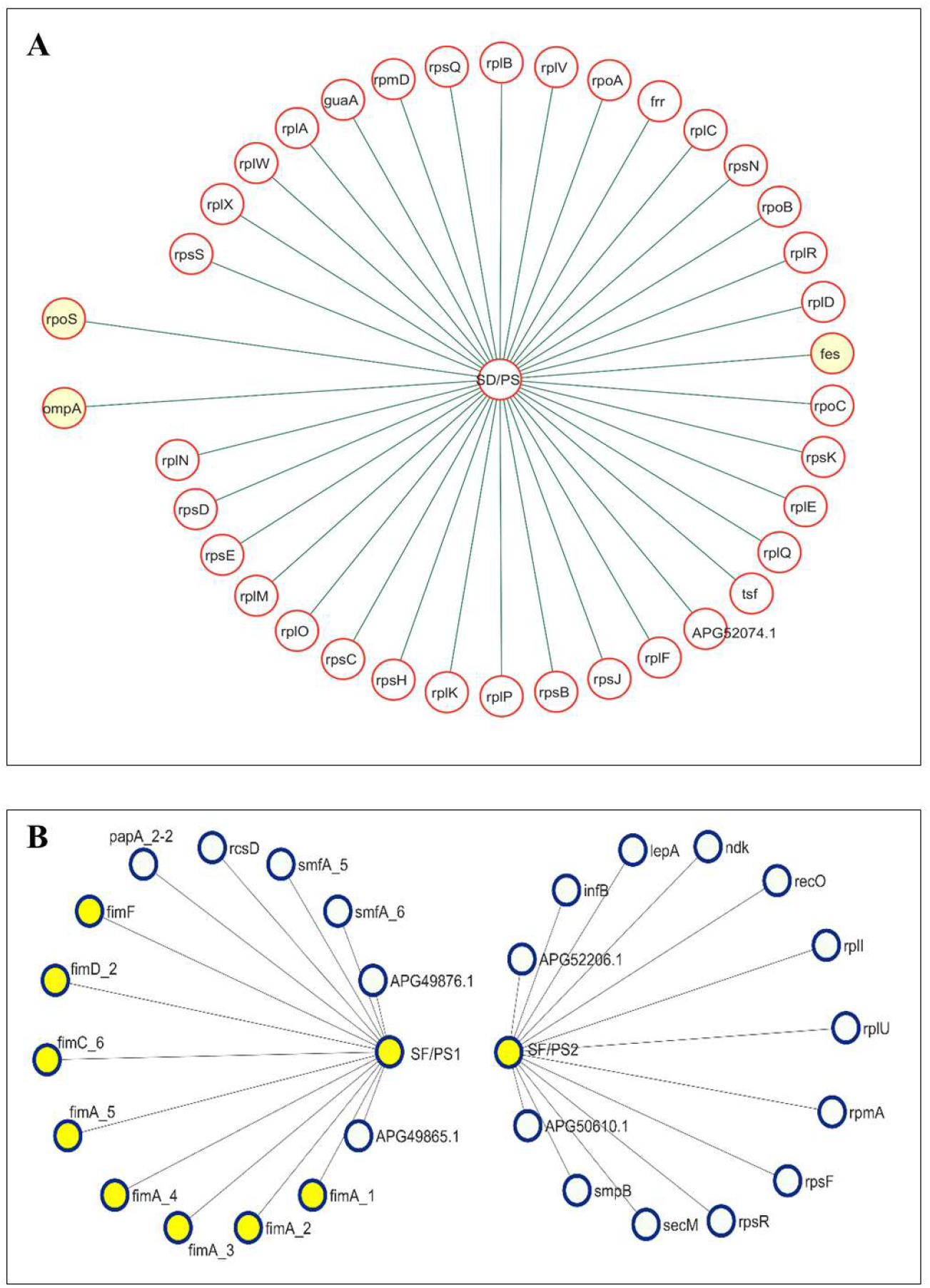
Microbial interaction of *Shigella dysenteriae* and *Shigella flexneri* virulence genes with *P. stuartii*. SD/PS1 and SD/PS2 cluster name; Red node: proteins of *P. stuartii*; ompA, fes and rpoS are virulence proteins of *Shigella dysenteriae* interacting with 27 ribosomal proteins of *P. stuartii* (**A**). Microbial interaction of *Shigella flexneri* virulence genes with *P. stuartii*. SP/PS1 and SP/PS2 are two cluster names; especially interacting with fimbrial virulence protein group of *S. flexneri*; Blue nodes: Proteins of *P. stuartii*; Yellow nodes: virulence proteins of *Shigella flexneri*. Solid lines are the interacting edges of the networks (**B**).

*Shigellla flexneri*, SF/PS1 cluster formed between the *P. stuartii* comprising 29 nodes and 27 edges containing fimbrial virulence protein groups. Specially, the Type-1 fimbrial protein A chain precursor fimA_1, fimA_2, fimA_3, fimA_4, fimA_5, fimD_2 and papA_2-2 was identified along with regulatory protein rcsD, metal ion transporter smfA_5, smfA_6 (**Figure 4B)**. SF/PS1 cluster was associated with the biological process of cell adhesion and molecular function of identical protein binding and subcellular localization of pilus. SF/PS2 cluster contained network of *Shigellla flexneri* with 14 nodes and 12 edges that contained only proteins of *Shigella flexneri* (**Figure 4B**).

*Salmonella enterica* Typhimurium established two distinct network clusters, ST/PS1 cluster comprised 17 nodes with 5 virulence genes (rpoS, fusA, sseA, ompA, and ratB) participating in network interactions with 14 proteins of *P. stuartii* namely polA_2, rpsA, rpmF, pheA, tyrA, smpB, csgD, pheT, rpsG, rpsL, rpsL, rplu, mgtB primarily ribosomal proteins, tRNA Synthetases and biofilm formation contributors. The ST/PS2 cluster consisted of 8 protein nodes and featured two known virulence genes, sseJ and InvF **(Figure 5A)**.

**Figure 5:**
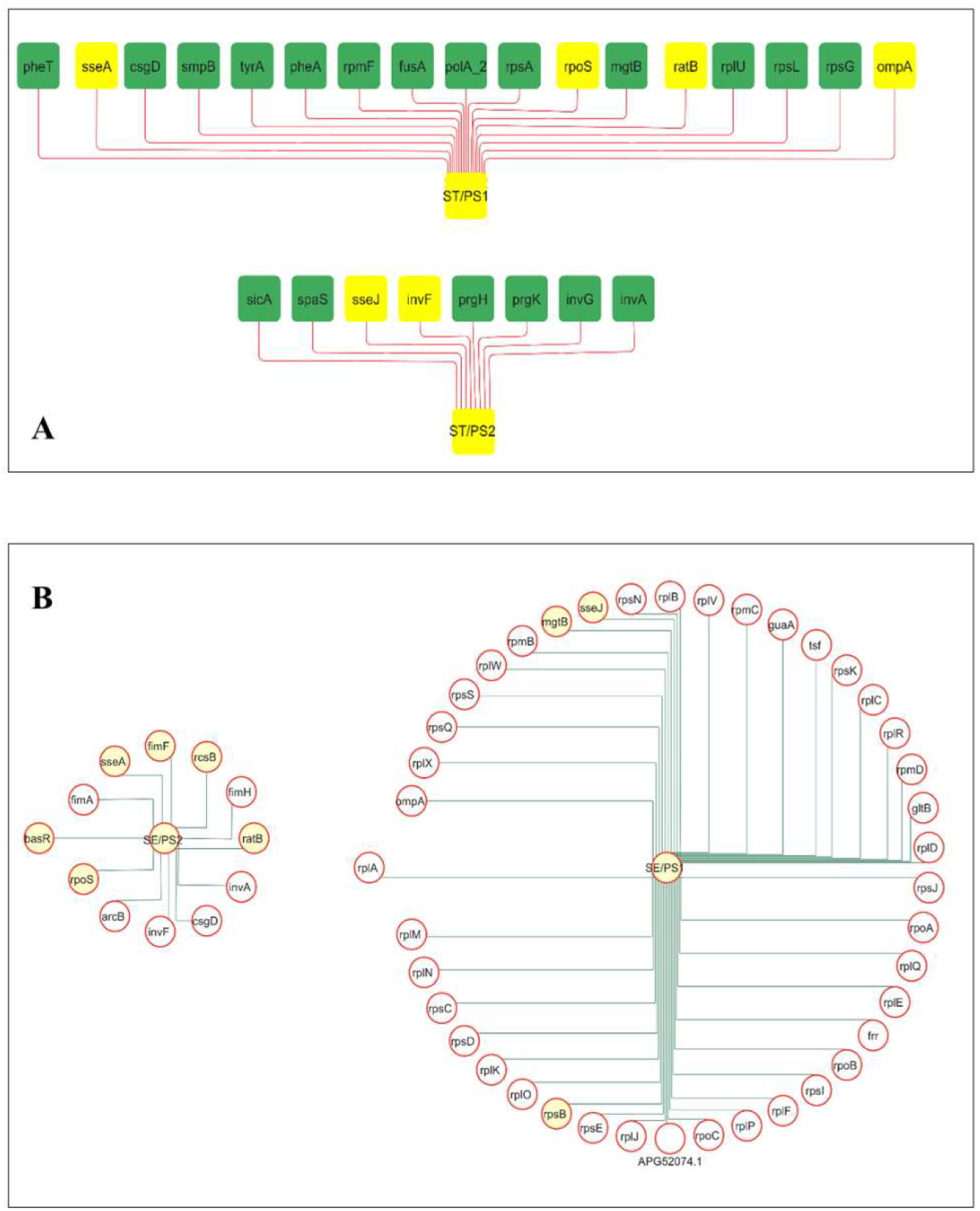
Microbial interaction of *Salmonella enterica* Typhimurium and *Salmonella enterica* Enteriditis virulence genes with *P. stuartii*. ST/PS1 and ST/PS2 are two cluster; Green nodes: Proteins of *P. stuartii*; Yellow nodes: virulence proteins of *Salmonella enterica* Typhimurium; five 5 virulence genes rpoS, fusA, sseA, ompA, and ratB involved in interaction with 14 *P. stuartii* proteins (**A**). *Salmonella enterica* Enteriditis virulence genes displayed the association with 36 *P. stuartii* proteins notably ribosomal protein groups and polymerase subunits. SE/PS1 and SE/PS2 are two clusters of *Salmonella* Enteriditis; Red nodes: Essential proteins of *P. stuartii*; Yellow nodes: virulence proteins of *Salmonella* Enteriditis (**B**). Solid lines are the interacting edges of the networks.

The ST/PS1 cluster’s biological processes mainly translation and aromatic amino acid biosynthesis, with molecular functions including chorismate mutase activity, RNA binding, and participation as ribosomal constituents. In contrast, the ST/PS2 cluster was situated in the Type III protein secretion system complex, emphasizing functions related to protein secretion.

SE/PS1 cluster of *Salmonella enterica* Enteriditis includes the virulence genes ompA, mgtB, sseJ, and rspB involving 36 proteins of *P. stuartii* including ribosomal proteins rplQ, rplE, rplF, rplR, rpl, rpsK, rpmD, rplP, rplJ, rplD, rplv, ffr, rpsC, rpsS, rplX, rplB, rplM, rplN, rplA, rpmB, rpsQ, rpsE, rplW, rplC, rplK, rplO, rpmC and rpsN. Three RNA polymerase subunits proteins rpoA, rpoB, rpoC and four other proteins guaA, gltB, tsf and APG2501.4 was visualized in the SE/PS1 cluster **(Figure 5B**). SE/PS2 is relatively smaller comprising 12 nodes and containing proteins such as fimA, fimF, invA, basR, rpoS, and ratB.

This cluster forms connections between virulence factors with proteins from *Providencia stuartii* including invF, sseA, arcb, rcsB, fimH, and csgD demonstrating interactions with this cluster **(Figure 5B)**. The SE/PS1 cluster’s predicted biological functions encompass translation, gene expression, and molecular functions related to ribosome constituents and rRNA binding also associated with RNA polymerase and the ribosome pathway. Conversely, the SE/PS2 cluster is presumed to be involved in the cellular component of the pilus and is associated with the family of fimbria proteins.

VC/PS1 cluster of *Vibrio cholerae* one virulence factor ompA that interacts with thirty *P. stuartii* proteins, primarily associated with biological processes such as translation, transcription, and gene expression. Molecular functions within this cluster include rRNA binding and serving as a structural constituent of the ribosome **(Figure 6A)**. On the other hand, cluster VC/PS2 represents 10 different nodes of virulence proteins interacting with *P. stuartii* proteins including the flg group, flagellar motor proteins like motB, chemotaxis response regulator protein cheW, and the two-component system response regulator, chemotaxis regulator cheY protein **(Figure 6A)**. Biological processes related to bacterial-type flagellar assembly, organization, cell motility, and chemotaxis. Additionally, the cluster exhibits strong associations with pathways involved in flagellar assembly, chemotaxis, and the two-component regulatory system.

**Figure 6:**
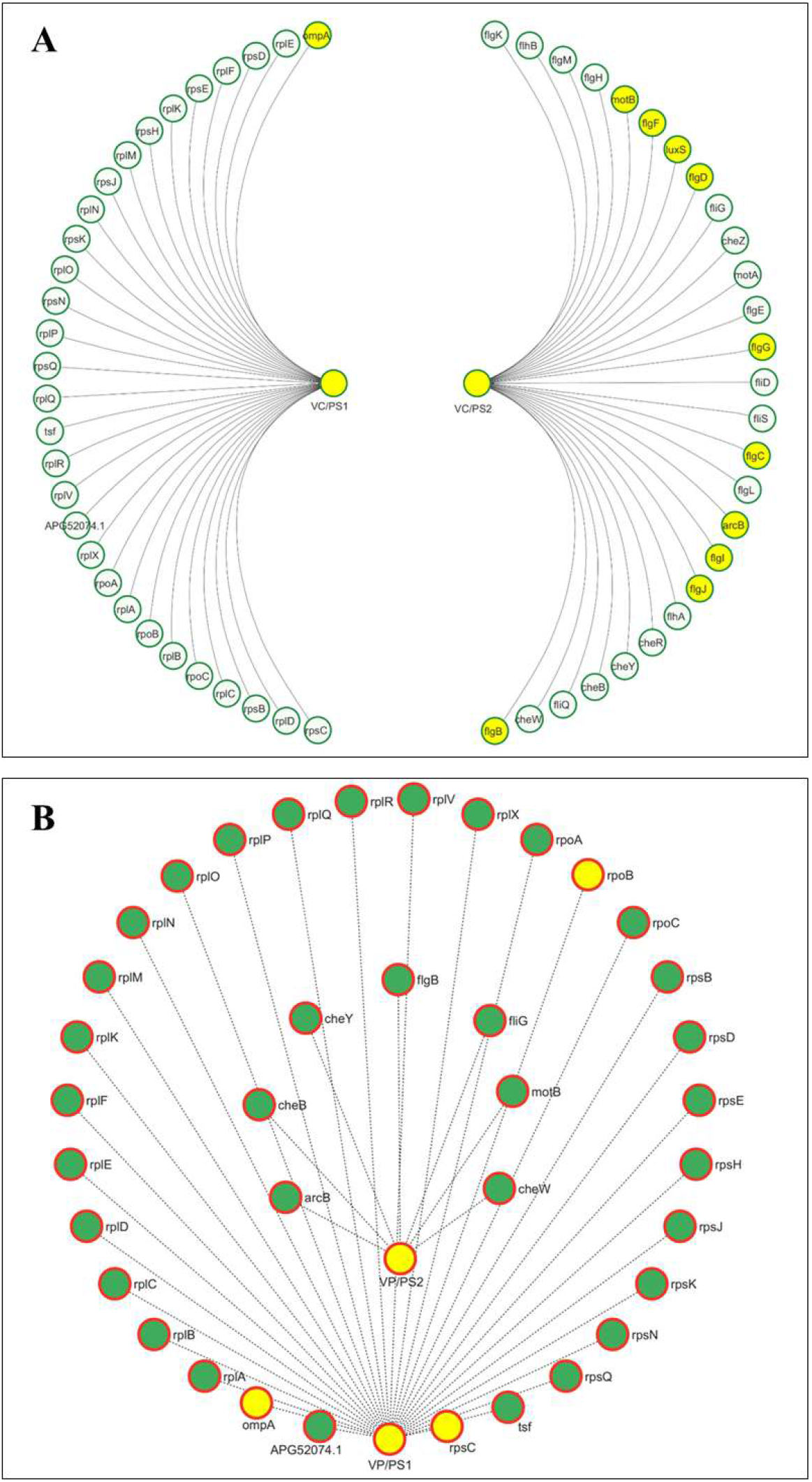
Microbial interaction of *Vibrio Cholerae* and *Vibrio parahemolyticus* virulence genes with *P. stuartii*. VC/PS2 cluster encompasses 10 virulence proteins interacting with P. stuartii flg group, motor proteins like motB, chemotaxis regulator protein cheW, and the two-component system response regulator, chemotaxis regulator cheY proteins; on the other VC/PS1 cluster have onE significant vir protein and the clusters is associated with translation, transcription, and gene expression. Green nodes: Proteins of *P. stuartii*; Yellow nodes: virulence proteins of *Vibrio Cholerae*. (**A**). VP/PS1 engages with the virulence gene ompA, rpoB and rpsC with 26 proteins of P. stuartii primarily ribosomal proteins and VP/PS2 VP/PS2 represents interactions among virulence genes encompassing cheB, cheY, cheW, flgB, fliG, and motB; Green nodes: Proteins of *P. stuartii*; Yellow nodes: virulence proteins of *Vibrio parahemolyticus*. Solid lines are the interacting edges of the networks (**B**).

*V. parahemolyticus* member of the *Vibrio* group, also exhibits interactions through the formation of two distinct protein clusters. The first cluster, VP/PS1 engages with the virulence gene ompA, rpoB and rpsC with 26 proteins of *P. stuartii* primarily ribosomal proteins. This cluster is associated with predicted biological functions such as transcription, translation, and gene expression also molecular functions involving rRNA binding and structural constituent of ribosome **(Figure 6B)**. The second cluster VP/PS2 represents interactions among virulence genes encompassing cheB, cheY, cheW, flgB, fliG, and motB, with one protein linked from *P. stuartii*, specifically the aerobic respiration two-component sensor histidine kinase A (arcB). This cluster is primarily associated with biological functions related to chemotaxis and signal transduction within the signal transduction pathway (**Figure 6B**).

The single network cluster formed between *C. jejuni* and *P. stuartii* was observed, CJ/PS cluster node proteins ompA, cheW, fliY and arcB represent the virulence factors of *C. jejuni* and they interact with 52 proteins from *P. stuartii* **(Figure 7)**. A significant single cluster of proteins interacted with the proteins majorly with ribosomal protein namely fabR, pstB, parC, elbB, aer, bfr, hemN_2, pstB, degQ, yidD, gltB, ysA_2-2, fliY, cysH, surA, trxA_2, hemE, ginL, syd, ppc, uvry, degS, pbpD_2, yycF, thiH, kdsD, dnaN, APG52436.1, uvrC, envZ, boxD, IptD, sthA, cheR, IptC, yihl, tar_2, rosB_1-2, glpG, thiE, hmp_2, ygdG, gppA, clpV1, caiD, cpxA and ginE. Additionally, twelve transport proteins i.e., pstB, zntR_1, kbl, envZ, boxD, IptD, sthA, cheR, IptC, tar_2, glpG, cpxA also other groups of enzyme and metabolic proteins namely fabR, hemN_2, gltB, kbl, ysA_2-2, ppc, degS, hemE, ginL, syd, dpiA, uvry, yycF, thiH, kdsD, dnaN, APG52436.1, uvrC, hmp_2, ygdG, gppA, clpV1, caiD, cpxA and ginE was involved in the network. Finally, 26 regulatory proteins and cell wall and membrane proteins of *P. stuartii*; degQ, syd, dpiA, uvry, degS, envZ, boxD, IptD, sthA, cheR, IptC, yihl, tar_2, rosB_1-2, clpV1, cpxA, parC, bfr, ppc, pbpD_2, yycF, galE_3, dnaN, APG52436.1, uvrC, ginE proteins linked with four virulence factors (**Figure 7**). This cluster is involved in biological processes related to chemotaxis, signal transduction, and membrane organization. Its molecular functions are associated with transmembrane signaling receptor activity, phosphorelay sensor kinase activity, and protein kinase activity, and plays a role in bacterial chemotaxis and the two-component regulatory system pathway.

**Figure 7:**
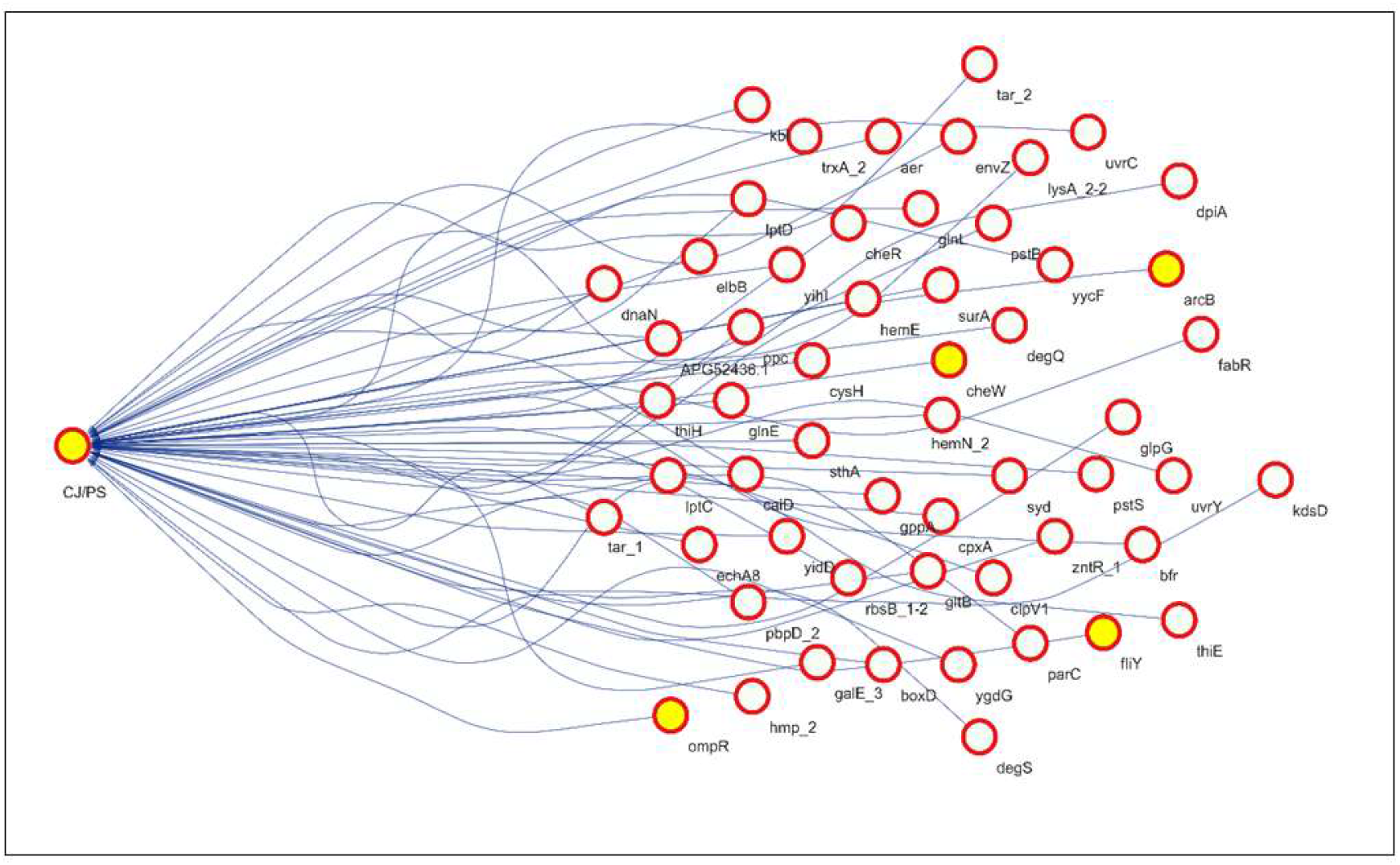
Microbial interaction of *Campylobacter jejuni* virulence genes with *P. stuartii*. CJ/PS cluster node proteins ompA, cheW, fliY and arcB represent the virulence factors of *C. jejuni* and they interact with 52 proteins from *P. stuartii*; Red circle nodes: Essential proteins of *P. stuartii*; Red circle yellow nodes: virulence proteins of Campylobacter jejuni. Curved lines are the interacting edges of this network.

Enteroinvasive *E. coli* formed the protein-protein interaction with 31 nodes and 29 edges. EIEC/PS1 cluster virulence genes are ompA and fus that interacts with 13 proteins of *P. stuartii* including rpsF, rplU, infB. hflB, eno, aceF, APG52206.1, Iucc, rpll, purA, pnp, mdh, and iutA associated with ribosomal, metabolism and protein synthesis **(Figure 8A)**. In this network, the biological processes (Gene Ontology) involve gene expression, peptide biosynthesis, and translation, while the molecular functions encompass rRNA binding and serving as structural constituents of ribosomes. On the contrary, EIEC/PS2 cluster encompasses rscB, fur and acrB virulence factors whereas the transcriptional regulator rcsB and ferric iron uptake transcriptional regulator is regulation factors and the more virulent acriflavine resistance protein (acrB) associated with the nuoC, ftsZ, ftsW, ftsQ, ftsl_1, rsmH, ftsA, dsbE, ddl, APG50799.1 and ftsL proteins of *P. stuartii* **(Figure 8A)**. This specific cluster is responsible for the cell shape regulation, cytokinesis and cell septum assembly and subcellularly localized in cell division site.

**Figure 8:**
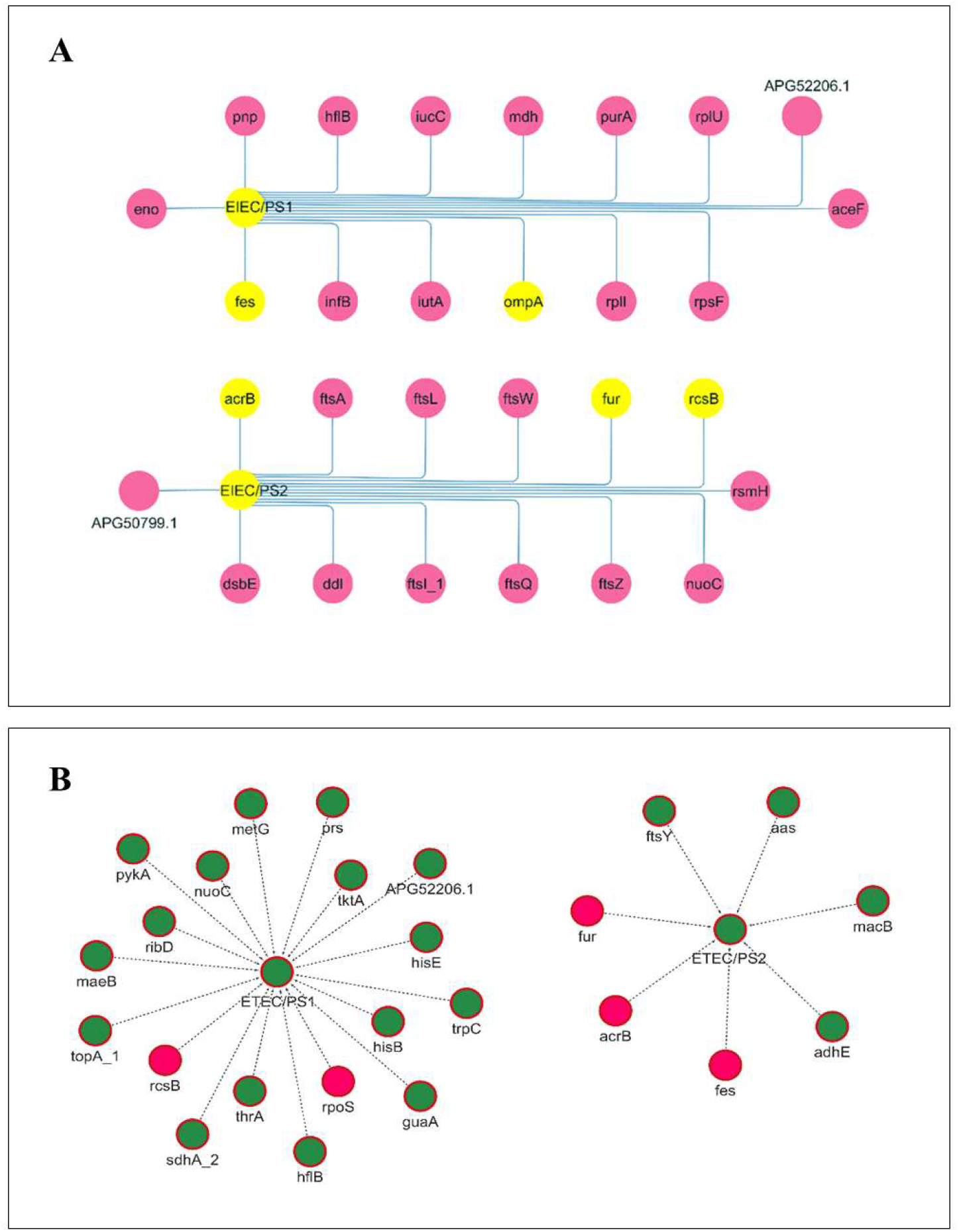
Microbial interaction of EIEC virulence genes with *P. stuartii*. EIEC/PS1 cluster have13 proteins of *P. stuartii* including rpsF, rplU, infB. hflB, eno, aceF, APG52206.1, Iucc, rpll, purA, pnp, mdh, and iutA associated with ribosomal, metabolism and protein synthesis whereas EIEC/PS2 cluster virulence factors fur, fes, and acrB along with four *P. stuartii* proteins associated with fatty acid degradation; Pink nodes: Essential proteins of *P. stuartii*; Yellow nodes: virulence proteins of Enteroinvasive *Escherichia coli*. Solid blue lines are the interacting edges of this network (**A**). ETEC/PS1 and ETEC/PS2 are two cluster names; Green nodes: Proteins of *P. stuartii*; Red nodes: virulence proteins of Enterotoxigenic *Escherichia coli*. Dotted lines are the interacting edges of this network (**B**).

The ETEC, a member of the *Escherichia Coli* pathogen family, ETEC/PS1 includes two frequently found virulence factors: transcriptional regulator (rcsB) and RNA polymerase sigma factor (rpoS) both involved in regulation processes with 16 proteins of the *P. stuartii* involves the metabolism and enzyme associated 16 proteins i.e., maeB, metG, prs, pykA, nuoC, APG52206.1, tktA, ribD, hisE, trpC, hisB, topA_1, thrA, guaA, sdhA_2, and hflB **(Figure 8B)**. This cluster’s biological process revolves cellular metabolic processes, role in histidine biosynthesis and the amino acid biosynthesis pathway. The second cluster, ETEC/PS2 comprises virulence factors fur, fes, and acrB along with four *P. stuartii* proteins associated with fatty acid degradation **(Figure 8B)**. Interestingly, virulence proteins of the bacterial pathogen *Clostridium difficile* have not exhibited any interactions with the opportunistic pathogen *P. stuartii*.

## 5. Discussion

Research on opportunistic bacteria and their interactions with diarrheal disease is limited in worldwide. Consequently, we carried out the investigation, which focused on opportunistic bacteria associated with diarrheal patients, particularly children diarrhea. In this regard, a clinical stool sample from a patient with diarrhea who was younger than ten years old was obtained, cultured, and microscopic visualization verified the existence of *Providencia spp*. (**Figure 1**) as the yellow colonies with short rods were appeared. The small, yellow rods confirm that this is the *Providencia spp*. [43]. Furthermore, *P. stuartii* was recognized as the presence of bacteria using phylogenetic analysis and 16srRNA sequencing (**Figure 2**). The improved comprehension of the genetic factors involved in human diseases, along with the rapid progress in sequencing technology, is revolutionizing the efficiency of diagnosing patients and offering greater possibilities for personalized treatment strategies [44]. Therefore, complete genome sequencing of *P. stuartii* bacteria was performed with the aid of Next Generation Sequencing technology (NGS) which has 4011 coding sequence among and the 1353 hypothetical proteins (**Figure 3**). Genome mapping has the potential to enhance our comprehension of genetic characteristics. Subsequently, we endeavored to examine and depict the microbial interplay among prominent bacteria responsible for causing diarrhea, leading to undertaking the compilation and annotation of 100 whole genome sequences (WGS) including *Shigella, Salmonella, Vibrio, Campylobacter, Clostridium,* and *Escherichia* species. In total, the coding sequences of the 10 bacterial strains encompassed a total of 3,28,714 protein sequences, which includes both essential and hypothetical proteins. Following the assembly and annotation process, our investigation focused on identifying the virulence genes of the diarrheal bacteria. These genes have been previously associated with the virulence of bacteria, specifically in relation to protein sequences or genes that induce acute or chronic diarrhea in both children and adults [45]. A specific group of genes will serve as crucial contributors to the bacterium’s capacity to induce pathogenicity. The gene products that enable the bacteria to colonize and survive within the host, or induce impairment to the host, are recognized as determinants of virulence or pathogenicity [45]. Our analysis was conducted in conjunction with the VFDB database. The 100 WGS of 10 most common pathogenic diarrheal bacteria yielded 534 virulence genes which are thought to be essential to cause the diarrhea.

The genetic composition of microorganisms governs their physiological functions, which are, in turn, modulated by the prevailing environmental factors [46]. It is noteworthy that microbes engage in interactions with one another to perform specific functions, and investigations into these interactions have significant value in comprehending their biological activities [47]. The *P. stuartii* and 100 WGS of 10 bacteria were from the diarrheal stool samples, and the *P. stuartii* was then employed to all 4011 virulence genes of 10 bacteria to make a network using the systems biology approach for the check of their microbial interactions. Systems biology is an increasingly important field which has the potential to unravel various biological complications with the help of computational and mathematical models. Now, it is very clear that the network construction of various genes present in such systems is supportive in revealing community-level interactions. The finding suggests that the overall network of the *P. stuartii* encompasses 2267 nodes and 8238 edges connecting the total of 4011 proteins involving both essential and hypothetical (**Supplementary Figure S2**).

The 33 proteins of *P. stuartii* made an interaction with ompA, fes and rpoS virulence genes of *S. dysenteriae*, making a biological cluster which is responsible for translation and subcellular localization around the polysomal ribosomes (**Figure 4A**). Within the highly interacting protein network of *Providencia stuartii*, the *Shigella dysenteriae* cluster revealed the presence of 33 essential genes (**Figure 4A**). Another strain of *Shigella spp*., *S. flexneri* formed the SP/PS1 cluster with 12 essential proteins of *P. stuartii* (**Figure 4B**). *Shigella flexneri* virulence gene interacting with the *P. stuartii* suggests the association in the process of cell adhesion and molecular function, interacting with two clusters (**Figure 4B**). The highly connected nodes of *P. stuartii* also revealed interactions with *Salmonella* Typhimurium in two clustered networks, involving 14 proteins (**Figure 5A**). The first cluster biological processes mainly translation and aromatic amino acid biosynthesis, with molecular functions including chorismate mutase activity, RNA binding, and participation as ribosomal constituents. In contrast, the second cluster was situated in the Type III protein secretion system complex, emphasizing functions related to protein secretion.

*Salmonella* Enteriditis displayed a remarkable number of interactions of its virulence genes with *P. stuartii*, involving 36 proteins (**Figure 5B**). *Vibrio cholerae* (*V. cholerae*) is one of the most pathogenic bacteria in causing diarrhea which made the cluster with *P. stuartii*, belonging to the flagellar group proteins (**Figure 6A**). *V. parahemolyticus* cluster formed an interconnected network of essential proteins with the bacterial networks, but no significant virulence genes interacted with the exception of ompA gene (**Figure 6B**). In the analysis of *Campylobacter jejuni*, a significant single cluster of proteins interacted with the proteins totaling 52 genes linked with four virulence factors (**Figure 7**). These clusters play an important role in the function of RNA polymerase and the ribosome pathway.

Enteroinvasive *Escherichia coli* established cluster with 13 proteins while on the other hand, ETEC/PS2 cluster also formed a strong indicative network, with microbial interaction (**Figure 8A**). Enterotoxigenic *Escherichia coli*, as a pathogen of diarrhea, displayed cluster-based interactions with maeB, metG, prs, pykA, nuoC, APG52206.1, tktA, ribD, hisE, trpC, hisB, topA_1, thrA, guaA, sdhA_2, and hflB proteins in the ETEC/PS1 cluster with the bacteria. Finally, ETEC/PS2 cluster of the interacting pathogen included ftsY, aas, macB, and adhE proteins associated with the virulence genes of *E. coli*. Interestingly in our findings, we did not find any interaction between the *C. difficile* and *P. stuartii* (**Figure 8B**).

A total of 207 proteins of *P. stuartii* were found to interact with the diarrheal pathogens, indicating their significant role in facilitating interactions with the microorganisms. The interaction could possibly elucidate the correlation between the presence of this particular bacterium and its identification in the clinical stool sample obtained from a patient suffering from diarrhea. These findings show that opportunistic pathogens, despite their classification as opportunistic, can function as dominant microorganisms in the development of diarrhea through microbial interactions. Hence, we argue that *P. stuartii* is the bacterium responsible for the onset of diarrhea.

## 6. Data availability

Genome sequence and Sequence read archive (SRA) were deposited in GenBank and under the accessions SAMN37419322 and SRP461633 in the BioProject PRJNA1017985.

## 8. Author Contribution

Mohammad Uzzal Hossain, Md. Shahadat Hossain, A.B.Z Naimur Rahman, Shajib Dey: Conceptualization; data retrieval; sequence analysis; laboratory experiments; writing – original draft; Zeshan Mahmud Chowdhury, Arittra Bhattacharjee: writing – review and editing; Ishtiaque Ahammad: manuscript preparations; Abu Hashem, Keshob Chandra Das: supervision; validation; Chaman Ara Keya: supervision; validation; Md. Salimullah: project administration; validation; supervision.

